# Towards environmental DNA-based bioassessment of freshwater reservoirs with small volumes of water: robust molecular protocols

**DOI:** 10.1101/2021.11.21.469426

**Authors:** Rebecca K. Loh, Sujatha N. Kutty, Darren C. J. Yeo, Rudolf Meier

**Affiliations:** Department of Biological Sciences, National University of Singapore, Singapore; Tropical Marine Science Institute, National University of Singapore, Singapore; Lee Kong Chian Natural History Museum, National University of Singapore, Singapore; Museum für Naturkunde, Leibniz Institute for Evolution and Biodiversity Science, Center for Integrative Biodiversity Discovery, Berlin, Germany

**Keywords:** Environmental DNA, Bioassessment, Small water volumes, Universal metazoan COI primer, Freshwater reservoir, DNA metabarcoding

## Abstract

Bioassessment of freshwater quality via eDNA is rapidly developing into a powerful alternative to traditional methods involving collecting, sorting, and identifying macroinvertebrates based on morphology. Particularly attractive would be methods that can use remote-controlled boats for sampling because it would allow for cost-effective, and frequent monitoring at multiple sites. The latter will be particularly important for tropical reservoirs that require year-around surveillance. We here optimize molecular protocols for capturing reservoir-specific differences in metazoan communities based on small water volumes (15 mL). The optimization is based on samples from two freshwater reservoirs with very different water qualities (“reservoir signal”). Each reservoir was sampled at three sites (“biological replicates”). For each water sample, the DNA was extracted twice (“technical replicates”). We then tested how much DNA template (0.1 ng to 15 ng) and how many PCR cycles (25 or 35) minimized variance between technical replicates. We find that 15 mL is sufficient for capturing the reservoir signal regardless of sampling time, template amounts, or PCR cycle numbers. Indeed, extrapolation from our results suggests that <1 mL would be sufficient because only 17 of 59 metazoan mOTUs (mainly planktonic crustaceans and rotifers) detected with a 313bp COI minibarcode were shared. We find that the use of 35 PCR cycles significantly lowered the number of detected species and that template amounts <0.5 ng yielded somewhat higher variance between technical replicates. Despite extensive trials, the variance between technical replicates remained high (Bray-Curtis: 5–20%; Jaccard: 10–40%) and we predict that it will be difficult to reduce this variance further. However, the overall reservoir differences are so strong that all biological and technical replicates can be correctly assigned.

## 1. Introduction

The bioassessment of freshwater environments based on environmental DNA (eDNA) is an important application of metabarcoding. It is increasingly replacing traditional approaches, which involved the collection and identification of macroinvertebrates (Cordier et al., 2018; Czechowski et al., 2020; Deiner et al., 2021; Fediajevaite et al., 2021). These traditional methods are unattractive in that they are invasive and require costly manual specimen collection, sorting and identification using morphological tools. Furthermore, it has been shown that the results are affected by the quality of sorting (Ji et al., 2013; Porter & Hajibabaei, 2018; Stein et al., 2014) and that the data are negatively affected by lack of taxonomic resolution (Sweeney et al., 2011). DNA-based methods, especially metabarcoding of water eDNA, promises to overcome many of these shortcomings (Porter & Hajibabaei, 2018; Ruppert et al., 2019) and large consortia are forming that explore the use of eDNA for assessing the biodiversity of freshwater habitats in many countries (e.g., DNAqua-Net: Bohmann et al., 2021; Leese et al., 2016).

Low biomonitoring frequencies are common in temperate and subtropical regions with distinct seasons and low water temperatures (UK: River Invertebrate Prediction and Classification System (RIVPACS); Queensland: Australian River Assessment System (AusRivAS); New Zealand (Stark & Maxted, 2007); USA, Oregon (City of Portland, 2016)). However, the needs are different in the tropics with year-round high temperatures and high levels of biological activity. Here, more regular monitoring of water quality is particularly important and would benefit from the availability of simple sampling techniques that are robust, avoid expensive equipment, and eliminate the need for highly trained field personnel. This is why we have been pursuing bioassessment based on eDNA from very small volume of water (15 mL). Such volumes can be retrieved using radio-controlled vehicles without filtration (e.g., New Smart Water Assessment Network, NUSwan (Chitre, 2018)).

As with all eDNA studies, it is important to carefully calibrate the sampling and molecular techniques before a technique is used (Aylagas et al., 2016; Cowart et al., 2015; Deiner et al., 2015; Goldberg et al., 2015; Hinlo et al., 2017; Piggott, 2016; Turner et al., 2014; Williams et al., 2016; Zaiko et al., n.d.). The minimum expectation is that the methods should be able to distinguish the water from reservoirs with different water qualities. This is what we here call the “reservoir signal”. On the other hand, water samples from adjacent sites in the same reservoir should not yield very different signatures (“biological replicates”: Zhan et al., 2014; Zhou et al., 2011) unless there are good reasons to expect pronounced quality differences within reservoirs. Low variability between biological sites is an attractive property given that most water management authorities are only interested in the overall water quality in a reservoir and only want to be alerted to site differences if the differences are so pronounced that they require management. Lastly, it is the goal of all laboratory and bioinformatics methods to minimize the signal obtained from technical replicates (Macher et al., 2021; Shirazi et al., 2021; Troth et al., 2021; Yamamoto et al., 2017; Zhan et al., 2014; Zhou et al., 2011) such as DNA extracted from subdivided water samples. Unfortunately, establishing appropriate sampling and molecular methods that have these desirable properties is very difficult because a large number of variables have to be tested before the robustness of protocols can be ascertained. Furthermore, there is no guarantee that a protocol yielding promising results for one system would also perform well for other water bodies. Regardless of these challenges, many water eDNA metabarcoding workflows are now undergoing calibration and refinement (Cristescu & Hebert, 2018; Kumar et al., 2020; McGee et al., 2019; Ruppert et al., 2019; Y. Wang et al., 2021; Zinger et al., 2019).

Bioassessment based on eDNA involves the detection of key taxa that can indicate water quality but it remains unclear whether ethanol precipitation and/or water filtration differ is more suitable for detecting relevant taxa. What is known, however, is that ethanol precipitation retains both intracellular and free-DNA, while filtration loses free-DNA (Goldberg et al., 2015; Turner et al., 2014); i.e., the techniques should not be mixed (Deiner et al., 2015; Hinlo et al., 2017; Peixoto et al., 2020; Piggott, 2016; Raemy & Ursenbacher, 2018; Spens et al., 2017). Not surprisingly, multiple studies have also shown that filtration retains more DNA and that this increases detection rates for vertebrates (Raemy & Ursenbacher, 2018; Spens et al., 2017; Turner et al., 2014). But the overall detection rates for other Metazoa remain poorly understood and may have been affected by inconsistent sample storage and DNA extraction methods across studies (both influence results: Deiner et al., 2015; Eichmiller, Miller, et al., 2016; Hinlo et al., 2017; Piggott, 2016; Williams et al., 2016). The most rigorous study was conducted by Deiner and colleagues (2015) who tested both methods on equal volumes of lake and river water. They concluded that water filtration recovered more eukaryotic genera, families and orders, including chironomids that are particularly useful for bioassessment. Compared to filtration, ethanol precipitation is operationally challenging whenever there is a need for processing large volumes of water (Deiner et al., 2015; Hinlo et al., 2017; Lear et al., 2018). Ethanol precipitation is thus less popular than the DNA extraction from filtrates. We here nevertheless pursue ethanol precipitation because routine bioassessment may not need the signal from large water volumes and small volumes are more easily sampled. Furthermore, direct DNA extraction from small water volumes is more suitable for water bodies with high amounts of particulate matter (Euclide et al., 2021) or high abundances of planktonic algae that clog filters. Such high abundances may be particularly common in tropical reservoirs (see Clews et al., 2014; Izmailova and Rumyantsev, 2016; Low et al., 2010).

Currently, there is also no agreement on how to standardize PCR conditions and reagents in eDNA metabarcoding studies. Yet, different primers targeting different fragments of a gene may result in the detection of different taxa, due to differing fragment sizes, degradation rates, or primer biases (Aylagas et al., 2016; Cowart et al., 2015). For example, higher annealing temperatures preferentially amplify templates with the smallest number of base mismatches at the priming site (Piñol et al., 2015; Sipos et al., 2007). Increasing the number of PCR cycles will furthermore bias abundance ratios against the rare sequences and sequences with more mismatches on the priming site (Piñol et al., 2015; Polz & Cavanaugh, 1998). Similarly inconsistent across studies is how DNA template amounts are standardized. Some studies standardize the amount of template directly by measuring its molecular weight (i.e., in nanograms, e.g., Antich et al., 2021) while others only standardize the volumes/weight of the water sample that was processed (e.g., Gleason et al., 2021; Martins et al., 2021; Meyer et al., 2021), although the eDNA concentration may differ widely. In other cases, PCR template amounts are not at all standardized, because DNA dilution has to be used to overcome PCR inhibition (e.g., Lim et al., 2016). Overall, it remains unclear to what extent these different approaches affect the Molecular Operational Taxonomic Units (mOTU) numbers detected for the same water eDNA. The existing evidence is ambiguous. For example, it is known for soil studies that diluted DNA template can still yield similar bacterial species richness estimates and community structure between technical replicates (Wu et al. 2010), but Castle et al. (2018) found that dilution may result in significant differences in fungal diversity.

The effect of the number of PCR cycles on community characterization also remains incompletely understood. Increasing PCR cycles is a convenient solution for eDNA samples with low DNA concentration, but it can increase the formation of chimeric sequences (Ahn et al., 2012). Furthermore, experimental studies have shown that using different numbers of PCR cycles can change the proportion of the diversity that is captured (Krehenwinkel et al., 2018), and that higher number of PCR cycles will skew template-to-product ratios (Piñol et al., 2015). Modelling also suggest that increasing the number of PCR cycles reduces detected species richness (Kelly et al., 2019). Overall, the available evidence suggests that the number of PCR cycles should be tightly controlled and probably be kept low. What is less certain is whether more repeatable results will be obtained with a larger or smaller number of PCR cycles although it has been suggested that decreasing the number of PCR cycles should reduce “PCR stochasticity” (Best et al., 2015; Kebschull & Zador, 2015). However, as far as we know, this relationship is only supported by theoretical models (Best et al., 2015).

Here, we test how the amount of DNA template and number of PCR cycles affect the repeatability of detecting the reservoir signal consisting of metazoan communities detected in small water quantities. We sample two tropical reservoirs that are 20 km apart and have very different water qualities and biodiversity (Bedok Reservoir and Pandan Reservoir: (Clews et al., 2014; Liew et al., 2018; Lim et al., 2016). For both reservoirs, three sites were sampled (“biological replicates”). Water samples were collected close to the surface, because we previously showed that the metazoan diversity captured with surface and benthic waters from the same reservoirs were very similar (Lim et al., 2016). eDNA was obtained from small water volumes via ethanol precipitation before extracting the DNA. For each water sample, we extracted the DNA for two subsamples (“technical replicates”) that were subjected to the same range of PCR conditions. Samples were subjected to two treatments — varying number of PCR cycles and DNA template amount (i.e., PCR template). Metazoan communities were studied using a universal COI primer that targets metazoans. We here try to identify the set of molecular conditions that reduces variability between technical replicates while retaining reservoir-specific signals. We find that similar molecular conditions minimized the variability between biological and technical replicates. However, we also find that this variability remains high regardless of whether it was assessed using presence/absence (Jaccard distance) or read abundance data (Bray-Curtis dissimilarity).

## 2. Materials and Methods

### 2.1 Field techniques

Van Dorn horizontal water samplers were used to obtain water samples at 0.5 m below the water surface in Bedok and Pandan Reservoir (Figure 1). The samples were kept on ice in clean glass bottles (newly purchased, sent for autoclaving and UV-radiated for at least 30 minutes before use) and transported back to the laboratory. Different samplers were used for the two reservoirs and the samplers were washed with 10% bleach solution and rinsed with > 2 L of RO water. Before taking the first sample at a new site, the sampler was rinsed twice with 250 mL of autoclaved, reverse osmosis purified (RO) water. All sampling equipment were handled with clean latex gloves, which were changed between sites. Water from the second rinse was kept as negative control (“sampling negatives”). 1 L and 500 mL of reservoir water was collected per site in Bedok Reservoir and Pandan Reservoir respectively. It is important to note here that such high volumes were only needed for the standardization experiment whereas future bioassessment could utilize much smaller volumes, down to 15 mL of water. All water samples were split into smaller volumes of 15 mL and topped up with 33.5 mL of molecular grade absolute ethanol and 1.5 mL of 3M sodium acetate, for preservation and ethanol precipitation (see Ficetola et al., 2008). Ethanol was kept at -80 °C for at least 24 hrs before use. Samples were stored at -80 °C until DNA extractions. All samples were kept in zip lock bags and stored in a dedicated freezer.

**Figure 1:**
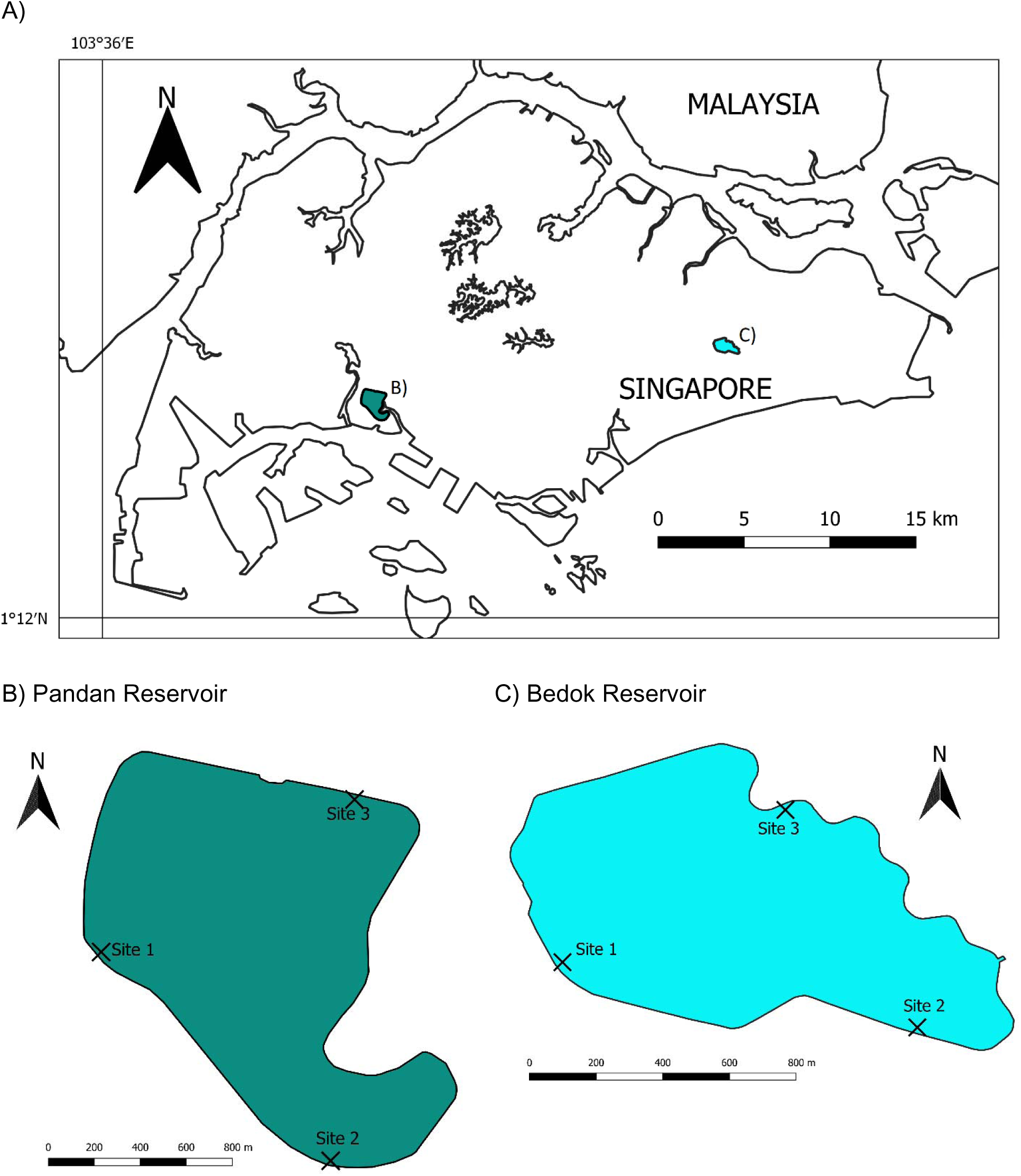
A) Map showing the two reservoirs sampled, in Singapore. Sampling sites at B) Pandan Reservoir and C) Bedok Reservoir, Singapore.

Both reservoirs were sampled in 2018, water samples were collected at three sites each (Figure 1. Refer to Table S1 in Supplementary Information for sampling dates and GPS coordinates).

### 2.2 DNA extraction

All benchwork was conducted in a class II biological safety cabinet. The working area was sterilized with 10% bleach and 98% methylated ethanol, followed by UV-radiation for at least 10 minutes. Two DNA extractions were performed for every water sample (“technical replicates”), resulting in 12 DNA extracts in total (Figure 2). For every DNA extract, 250 mL of reservoir water for Bedok samples and 100 mL of reservoir water for Pandan samples was extracted. More water needed to be DNA extracted from Bedok samples in order to obtain enough DNA for all experiments. DNA extraction utilized conventional phenol-chloroform protocol (see Supplementary Materials and Methods, Sambrook and Russell, 2006) although we have since established that bead-based automated extraction and commercial kits are also suitable. Negative controls were included in each round of DNA extraction (“extraction negatives”, nine in total), whereby empty Falcon tubes were used in place of sample tubes. Extracted DNA was dissolved in molecular grade water and kept at -30 °C until further processing. The amounts of eDNA (ng) obtained were quantified using a Qubit 2.0 Fluorometer (dsDNA HS Assay, Life Technologies) according to manufacturer’s protocol.

**Figure 2:**
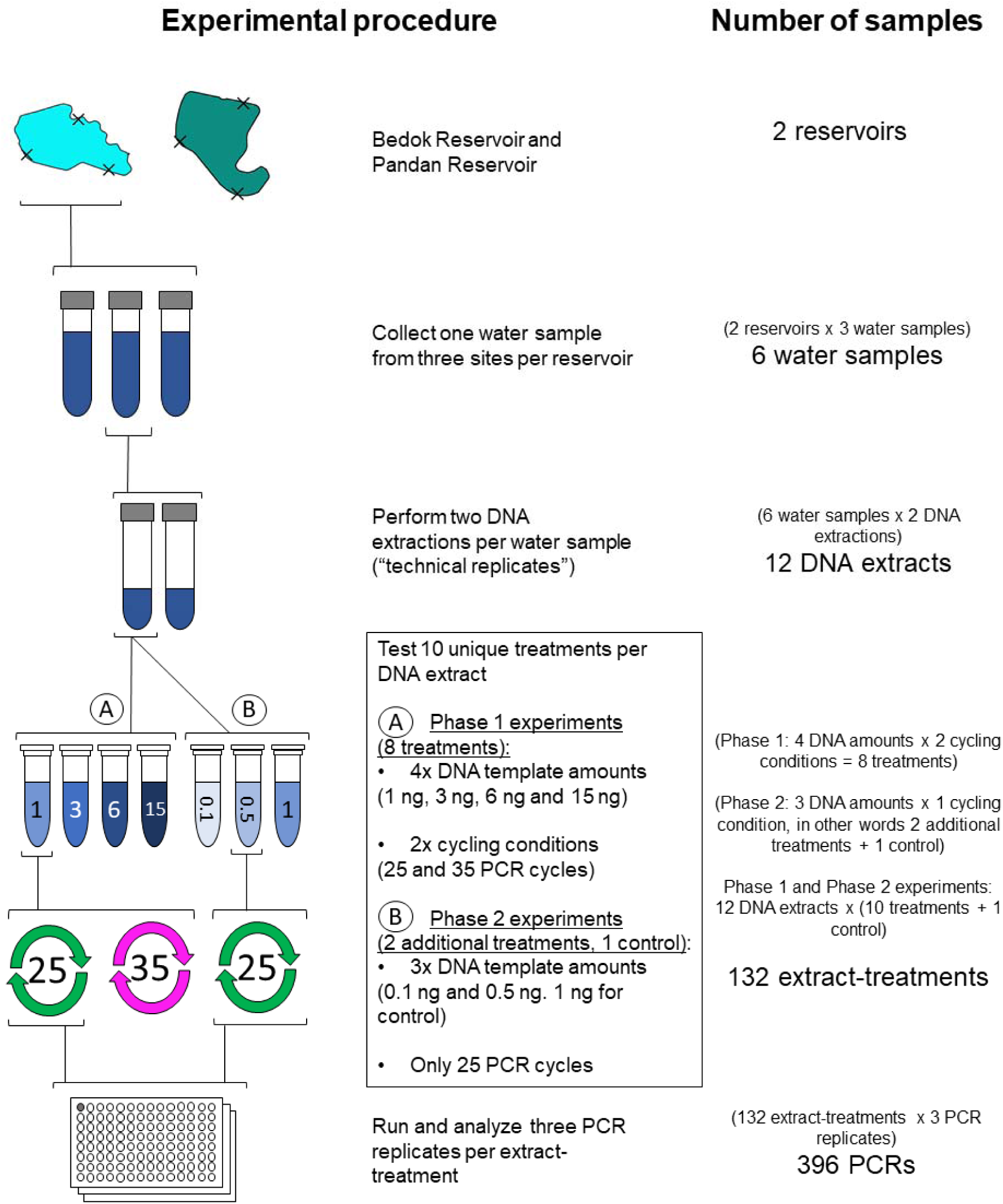
Summary of experimental design and procedure.

#### 2.3.1 PCR experiments

We varied the number of PCR cycles (25 or 35) and the amount of DNA template, and carried out laboratory experiments in two phases, approximately one year apart. In the first phase, DNA template was varied between 1 and 15 ng (4 treatments: 1 ng, 3 ng, 6 ng, 15 ng) and we tested 25 and 35 PCR cycles. The results indicated that 35 cycles reduced the number of robustly detected mOTUs (see below) and suggested that even lower amounts of template may be suitable as long as 25 PCR cycles were used. Therefore, in the second phase of laboratory experiments we used the same samples (i.e. 12 DNA extracts) to test lower template amounts (0.1 ng and 0.5 ng; and 1 ng again to use as control), but only used 25 PCR cycles in these trials. Note that these DNA amounts span the observed range of water eDNA that we have observed in 15 mL of raw water across 10 Singapore reservoirs (Loh, 2019).

Throughout the study we used the degenerate, universal metazoan primers mICOIintF: 5’-GGWACWGGWTGAACWGTWTAYCCYCC (Leray et al., 2013) and jgHCO2198: 5’-TAIACYTCIGGRTGICCRAARAAYCA (Geller et al., 2013) during PCR amplification. The primer targets a 313bp segment of the barcoding region of COI. Each PCR was performed with forward and reverse primers that have unique tags. All primer tags are 9 bp long and have at least 3 base-pair differences (Meier et al., 2016; W. Y. Wang et al., 2018). For every batch of PCRs (see Figure 2), negative controls were included that used molecular grade water in place of DNA template. PCR mixtures and cycling conditions are provided in Supplementary Materials and Methods. At least three PCR replicates were performed for each DNA extract and cycling condition (“extract-treatment”, see Figure 2). All PCR products were visualized using gel electrophoresis and failed PCRs were repeated with new tagged primers in order to control for the effect of freezing and thawing. The overall design of the project is depicted in Figure 2.

#### 2.3.2 Library preparation

All PCR products of the same treatment conditions were pooled in equal volumes and purified using a Serapure bead mixture prepared according to the Faircloth Serapure Protocol v2.3 (Faircloth & Glenn, 2014; Rohland & Reich, 2012). Purified PCR product pools were quantified using Qubit 2.0 and normalized between pools. DNA library preparation, using the TruSeq DNA PCR-free Library Prep Kits, and the addition of a 20% PhiX spike was carried out by Axil Scientific. Each library was sequenced by the Genomic Institute of Singapore on an Illumina HiSeq 2500 lane using the HiSeq Rapid SBS Kit v2 in Rapid Run Mode, for 250 bp paired end sequencing. To avoid batch effects (i.e., differences between PCR machines, PCR reagent batches or different DNA libraries), PCR replicates of the same extract-treatments were not processed in the same batch (i.e., different batch of reagents, processed in different 96- samples PCR batches and pooled in different DNA libraries).

### 2.4 Quality filtering and normalizing read coverage

We processed the metabarcoding sequence data using a mixture of well-established bioinformatic programs and in-house scripts (refer to Table 1 for detailed steps of the bioinformatic pipeline applied in chronological order). Briefly, forward and reserve reads were merged using Paired-End reAd mergeR (Zhang et al., 2014) and demultiplexed (assigned to PCR samples) using OBITools (Boyer et al., 2016). Metazoan sequences were identified using a BLAST search against sequences from NCBI GenBank and our internal reference library, using BLAST+ 2.2.30 (Camacho et al., 2009). Out internal reference library primarily contains curated COI DNA/NGS barcodes of macroinvertebrates collected from Singapore freshwater and mangrove habitats (NCBI GenBank, awaiting accession numbers). For each extract-treatment, we retained only the three PCR samples with the highest read coverage for further analyses. The read coverage of each PCR sample was then normalized to the same level — 22,002 reads per PCR sample, matching the lowest read coverage in the dataset — using SeqKit (Shen et al., 2016). Using OBITools (Boyer et al., 2016), low abundance sequences (<4 read counts) within each PCR replicate were discarded and sequence variants with only one base difference across the dataset were combined. Sequences containing stop codons in all translation frames (SeqKit) and unique sequences detected in only one out of three PCR samples of the same extract-treatment were discarded. The retained sequences were aligned using MAFFT v1 and poorly aligned sequences as visualized on AliView (Larsson, 2014) were excluded. Molecular Operational Taxonomic Units (mOTUs) were delimited using objective clustering (Meier et al., 2016) as implemented by a new script (Yeo et al., 2020) based on 3% pairwise-distances (p-distance). mOTUs detected in negative controls were removed from all samples.

**Table 1:**
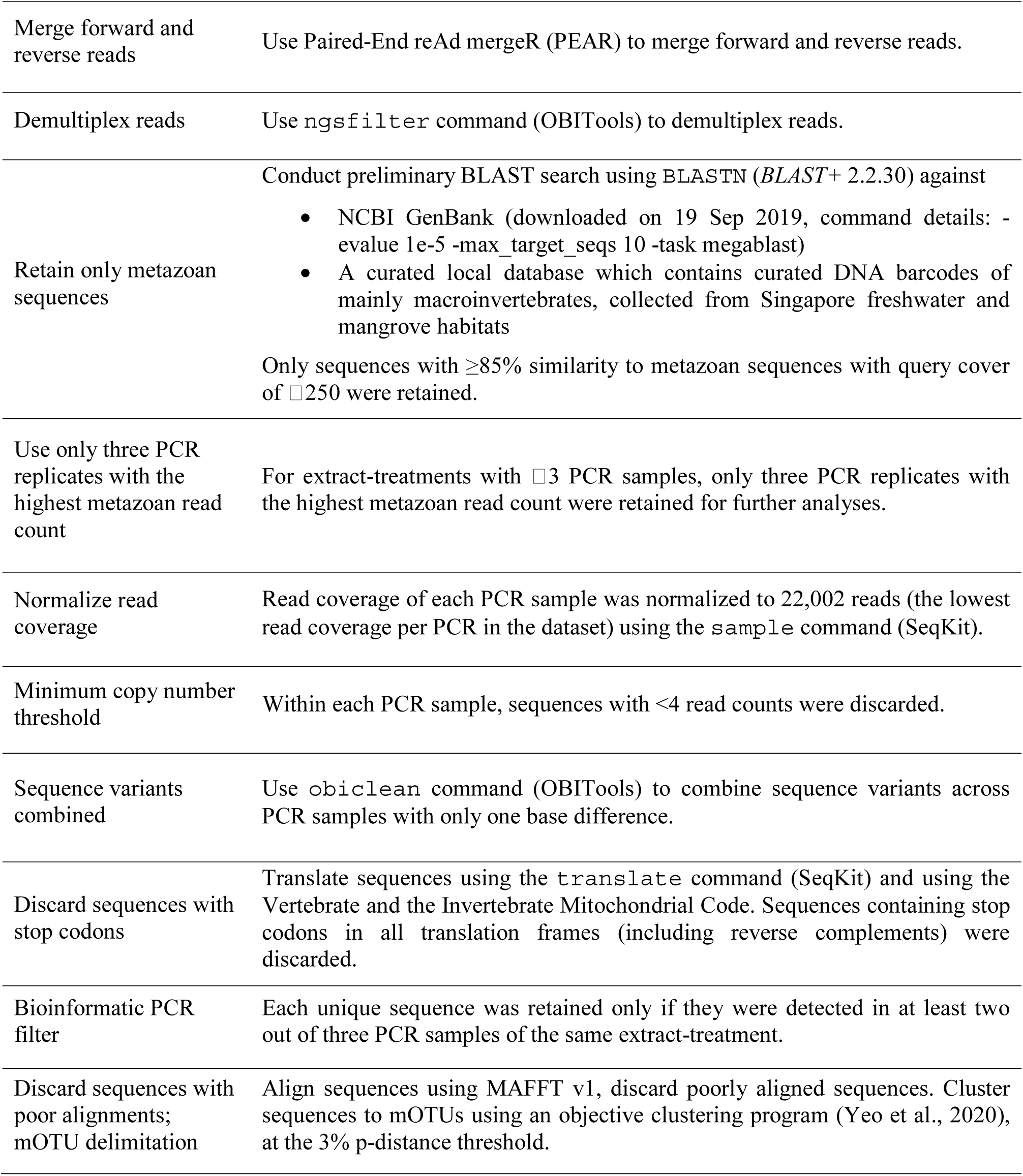
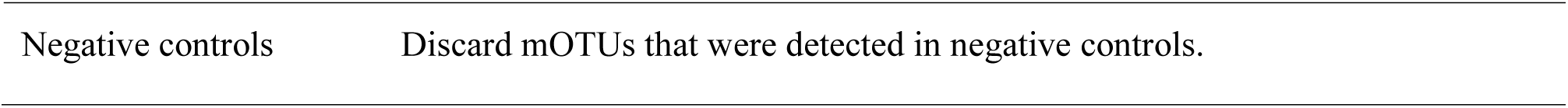
Bioinformatic pipeline.

### 2.5 Statistics analyses

All statistical analyses were performed in *R* v3.5.2. In order to quantify the repeatability of results, pairwise dissimilarity was calculated for the species compositions of technical replicates (i.e., DNA extracted from the same water sample and subjected to the same molecular conditions, N=132) and biological replicates. Note that for each set of molecular conditions, there are six possible pairs of biological replicates (3 sites × 2 DNA extracts) per reservoir. Hence, only the minimum and maximum values are reported. Two dissimilarity indices were used. The first was Jaccard distances, using binary (presence/absence) data for mOTUs. The second was Bray-Curtis dissimilarity, using relative read counts as a potential proxy for mOTU abundance.

To evaluate the significance of treatment effects on the repeatability of technical replicates, we used statistical modeling with model simplification as well as the information-theoretic approach. We constructed models using functions in the R package *lme4* v1.1-27 (Bates et al., 2015). The response variable is the pairwise dissimilarity between technical replicates (Jaccard distances or Bray-Curtis dissimilarity), which is an indication of variance; the explanatory variables are the treatments we are evaluating: the number of PCR cycles and the amount of DNA template used during PCR amplification. Since our response variable ranges from 0 to 1, we initially started with a binomial generalized linear mixed-effects model (GLMM) using the `glmer` function, and Laplace approximation (Bolker et al., 2009). This allowed us to include random effects to account for differences in the reservoir sampled. However, the model was overfitted, so we excluded random effects and refitted the data with a binomial generalized linear model (GLM) using the `glm` function. After fitting the maximal model, residuals were checked to ensure that there were no problems with overdispersion. The fixed effects of the maximal model includes the explanatory variables (“`cycles`”, “`DNA`”), a quadratic term for DNA template amounts (“`DNA`^2^”), as well as all possible interactions between terms. Starting with the maximal model, we simplified the model by discarding non-significant predictors (p-value<0.05) in a stepwise manner to reach the minimum adequate model. Treatment effects were further analyzed by using an information-theoretic approach. We generated a number of candidate models using different combinations of the explanatory variables, and compared using AICc, which corrects for small sample sizes (function `model.sel`, R package *MuMIn* v1.43.17 (Bartoń, 2020)). Refer to Supplementary Materials and Methods, “*List of candidate models assessed using the information-theoretic approach (1)”*, for the list of candidate models.

We also evaluated the effect of each treatment on the number of mOTUs detected (i.e., mOTU richness) using statistical modeling and the information-theoretic approach. The response variable is the number of mOTUs detected in a sample, or mOTU richness; the explanatory variables are the treatments we are evaluating; the random effect accounts for differences in the reservoir sampled. Since our response variable is count data, we fitted poisson GLMMs to data using the `glmer` function and Laplace approximation, and checked that the residuals showed no overdispersion (checked using `overdisp.glmer`, R package *RVAideMemoire* v0.9-80 (RRID:SCR_015657)). The fixed effects of the maximal model includes the explanatory variables (“`cycles`”, “`DNA`”), a quadratic term for DNA template amounts (“`DNA`^2^”), as well as all possible interactions between terms. The maximal model was simplified in a stepwise manner to reach the minimum adequate model. The amount of variance explained by the minimum adequate model was extracted using the `r.squaredglmm` function as implemented in the R package *MuMIn* (Bartoń, 2020). We also used the information-theoretic approach, by generating and comparing candidate models (AICc). Refer to Supplementary Materials and Methods, “*List of candidate models assessed using the information-theoretic approach (2)*”, for list of candidate models.

To compare mOTU compositions between multiple samples, pairwise dissimilarities were visualized on two-dimensional non-metric multidimensional scaling (NMDS) plots using the `metaMDS` function as implemented in *vegan* v2.5-7 (Oksanen et al., 2018).

## 3. Results

### 3.1 Sampling coverage, quality filtering and controls

In total, 167,814,405 reads were demultiplexed, and 351,322 ±163,937 and 496,225 ±217,401 read counts were assigned to Bedok and Pandan PCR samples respectively. Despite standardizing the amount of DNA template added during PCR amplification, Bedok samples still yielded lower proportion of metazoan read counts than Pandan samples (Figure 3). Hence, it was necessary to normalize metazoan read counts across PCR replicates.

**Figure 3:**
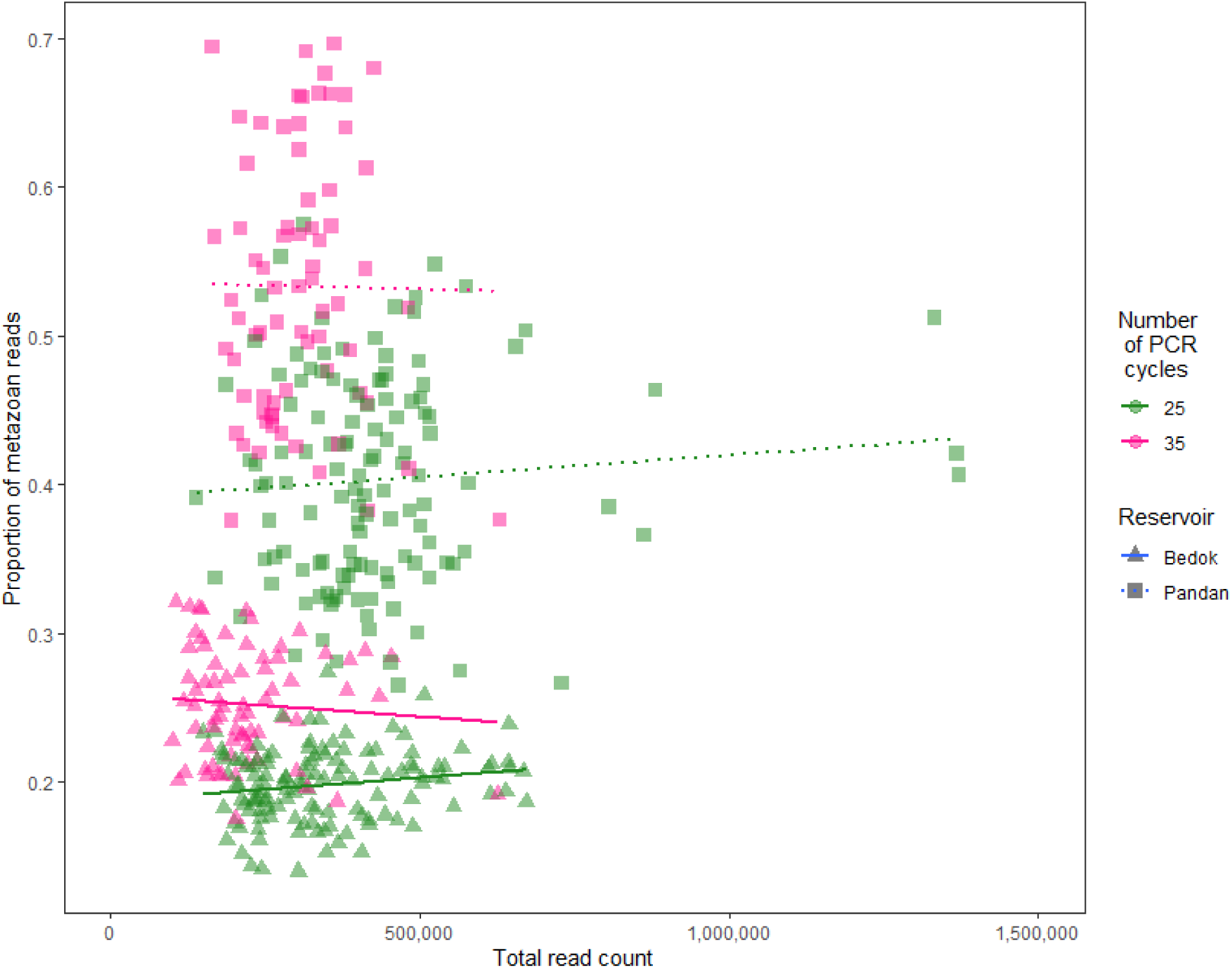
Pandan samples consistently yielded higher proportions of metazoan reads per PCR sample.

Although we conducted laboratory experiments in two phases, experimental controls (i.e., shared 1 ng treatment between phases) yielded similar results, which indicates the time of lab work was not a confounding factor. Hence, we combined the data from the different experiments. After applying all quality filters, only one mOTU was detected in the PCR negative controls (*Hypena obesalis* on Genbank: 94.3% similarity, and Lepidoptera sp. on BOLD Systems: 94.3% similarity) and was removed from all samples.

### 3.2 Reservoir-specific patterns

We observed similar numbers of mOTUs in the two reservoirs with very different water quality although the eDNA concentration of was considerably higher in Pandan (10.2±3.15 ng/mL) than Bedok Reservoir (2.40±0.169 ng/mL). A total of 59 mOTUs (3% p-distance threshold) satisfied all quality filters and the mOTU compositions detected showed clear reservoir-specific differences. Only 17 out of 59 mOTUs (28.8%) were found in both reservoirs (Table S2 in Supplementary Information). Most of the identifiable mOTUs were zooplankton and the reservoir-specific patterns were so strong that all water samples could be unambiguously assigned to reservoirs regardless of which molecular procedures were used — in other words, the eDNA in 15 mL of water is sufficient for distinguishing Bedok and Pandan reservoir water.

*3.3 The effects of PCR cycle numbers and DNA amount* The use of higher numbers of PCR cycles (35x) and very low DNA template amounts during PCR (<0.5 ng) depresses the number of mOTUs for both reservoirs. The change in species composition is due to losing mOTUs detected with low read abundances (Figure 4); i.e., fewer mOTUs were detected when 35 PCR cycles or 0.1 ng of DNA template was used (Figure 5), although the overall species composition remains similar as illustrated by NMDS plots (Figure 6). GLMM analysis showed that only the number of PCR cycles explained a statistically significant amount of variance in the model (Table 2, p<0.05). But comparing models using AICc indicated that at least one of the top two models with highest support (Δ 2: Table 3) contained PCR cycle numbers and DNA amount which suggests that the latter also explains the variation in mOTUs richness to some extent.

**Figure 4:**
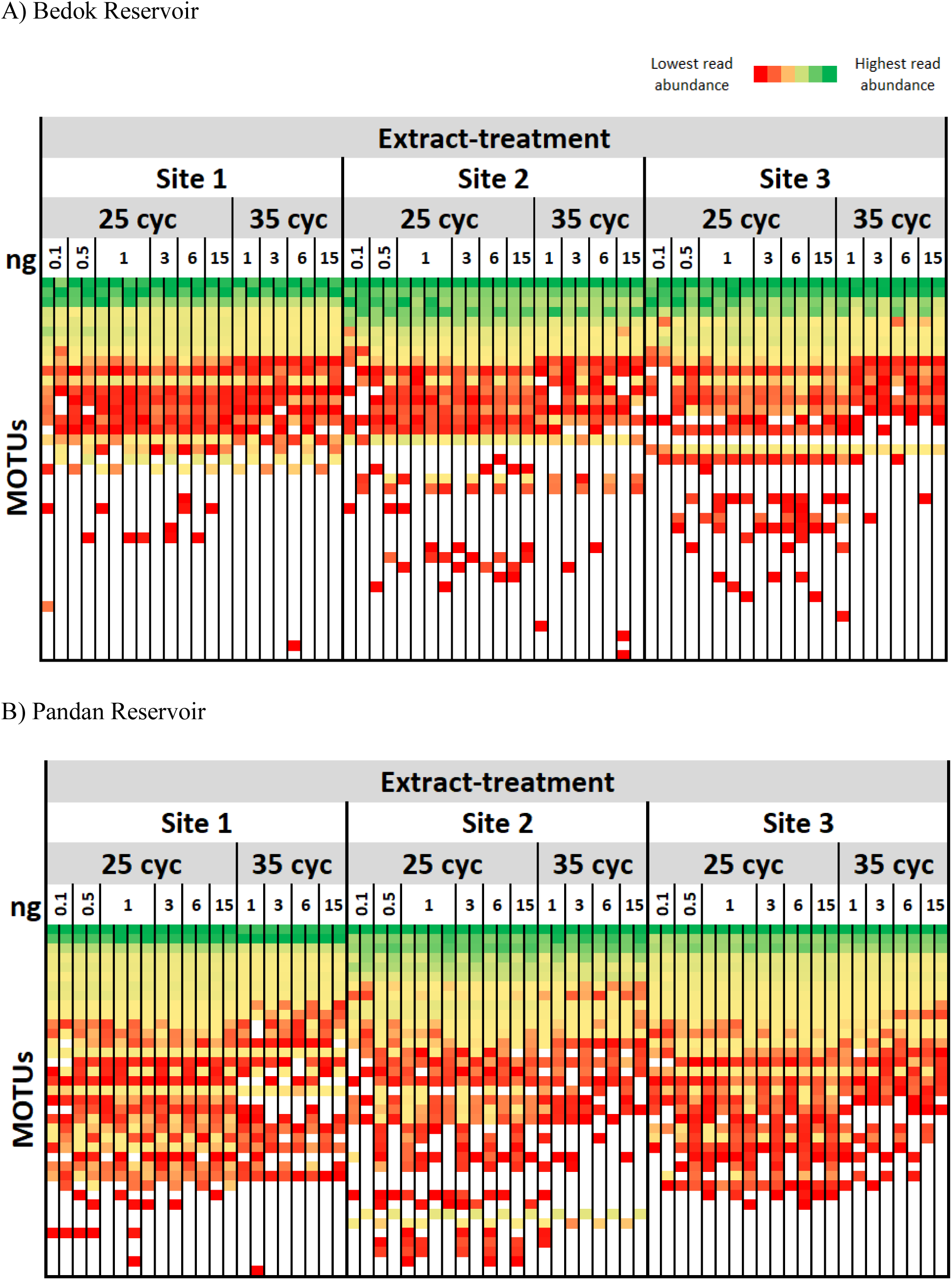
Heatmaps suggesting that low abundance mOTUs were less likely to be detected when 35 instead of 25 PCR cycles were employed, and when very low PCR template amounts (0.1 ng) were used. Colors (see legend) reflect the ranked read abundances of all mOTUs (different rows) within an extract-treatment (each column). mOTUs are arranged from top to bottom by total read abundances.

**Figure 5:**
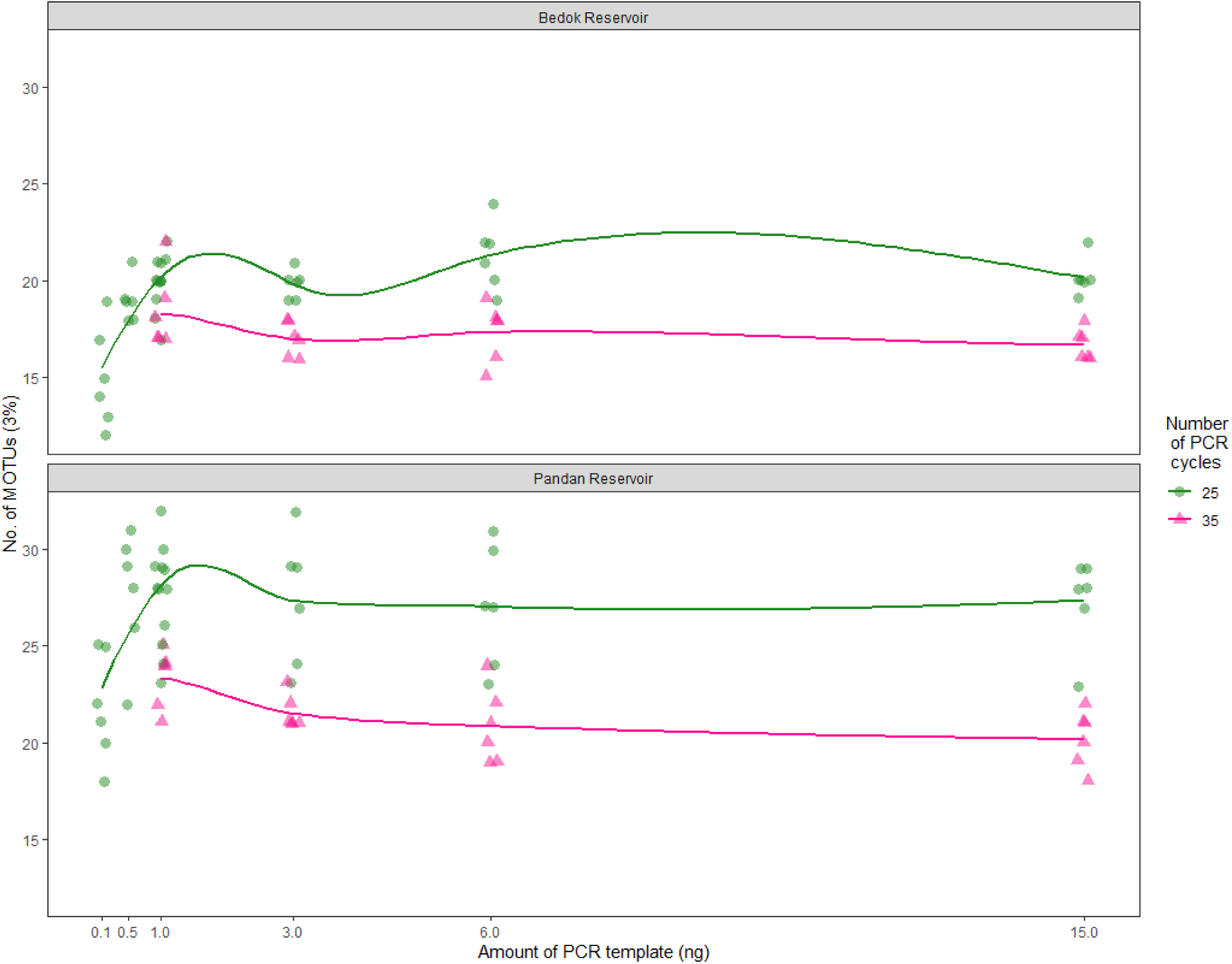
Lower numbers of PCR cycles consistently yielded higher numbers of mOTUs across all amounts of DNA templates tested. Each data point is an extract-treatment.

**Figure 6:**
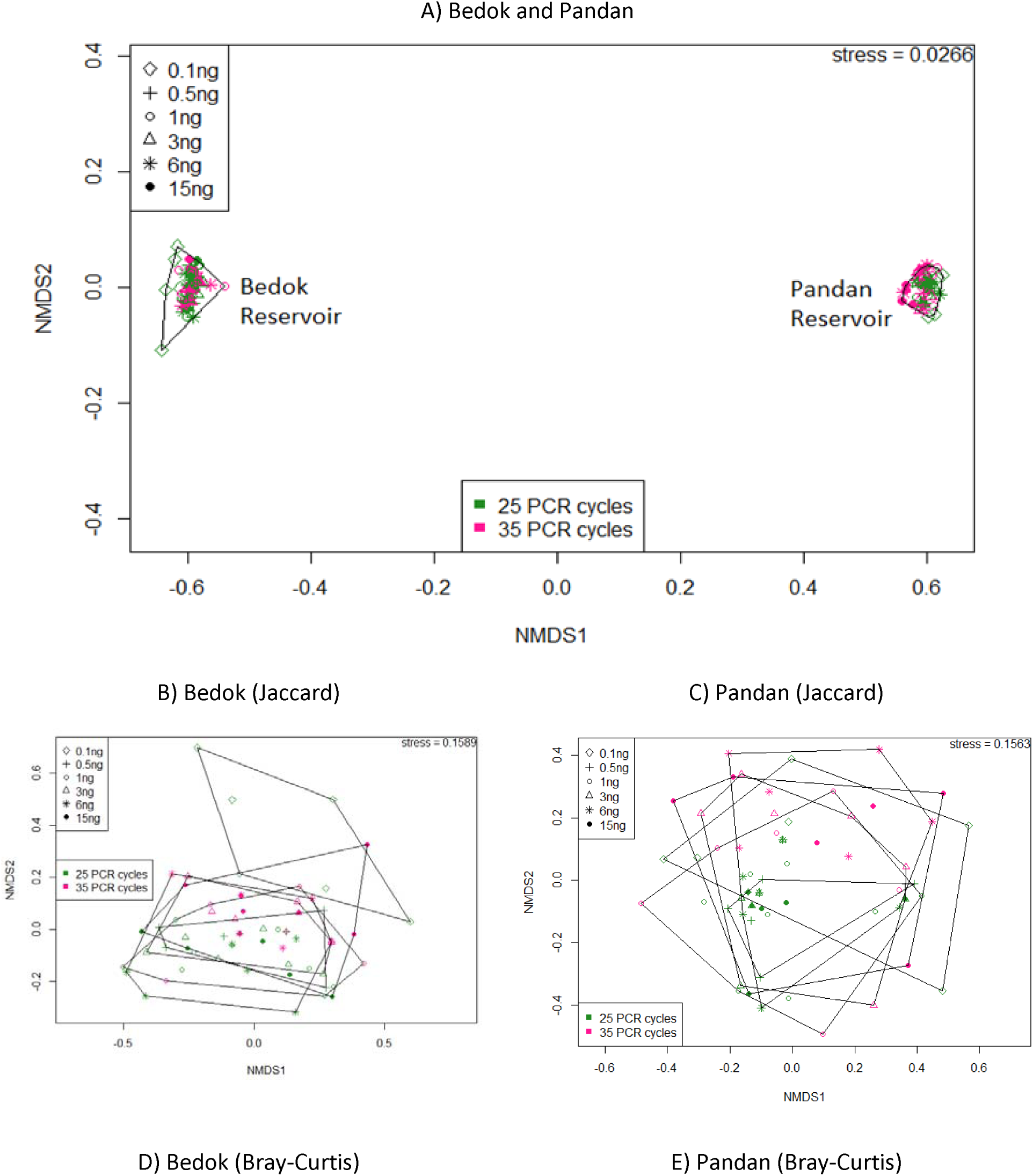
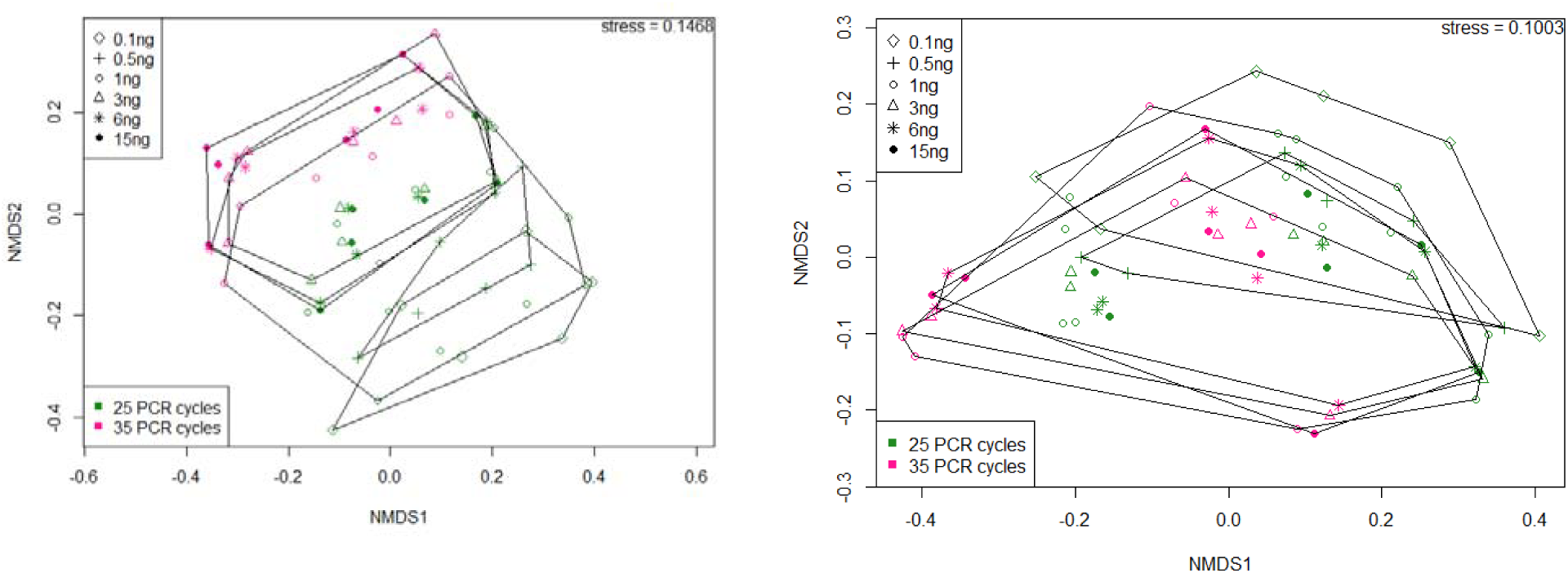
Differences in mOTU communities were primarily driven by reservoir differences instead of differences in the molecular protocol. A) NMDS plot of all extract-treatments (N=132) in the present study, samples are compared using Jaccard distances (presence/absence data). B) NMDS plots of Bedok samples and C) Pandan samples only, samples are compared using Jaccard distances (presence/absence data). D) NMDS plots of Bedok samples and E) Pandan samples only, samples are compared using Bray-Curtis dissimilarity (relative read counts).

**Table 2:**
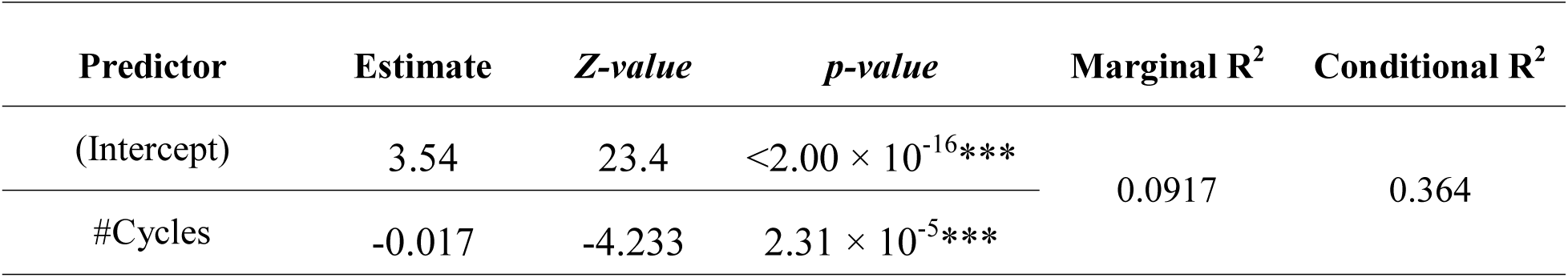
The minimum adequate model after stepwise model simplification of the maximal model (GLMM), showed that the number of PCR cycles (“#Cycles”) explained a significant amount of variation in mOTU richness.

**Table 3:** At least one of the most highly supported GLMMs (ΔAICc<2) showed that both variables: the number of PCR cycles (“#Cycles”) and the amount of DNA template (“DNA”), are important in explaining the variation in mOTU richness. Table shows the ΔAICc<2 and model weights (*w)* for the top six most supported models using the information-theoretic approach.

| Model Fixed Effects | $\Delta\text{AICc}$ | $w$ |
| --- | --- | --- |
| #Cycles | 0.00 | 0.354 |
| #Cycles + DNA + #Cycles:DNA | 1.49 | 0.168 |
| #Cycles + DNA | 2.04 | 0.127 |
| #Cycles + DNA<sup>2</sup> | 2.13 | 0.122 |
| #Cycles + DNA + DNA<sup>2</sup> | 2.56 | 0.098 |
| #Cycles + DNA<sup>2</sup> + #Cycles: DNA<sup>2</sup> | 2.65 | 0.094 |
| All other models | $\geq 4.58$ | $\leq 0.036$ |

When studying the relationship between cycling conditions, template amount and variance between technical replicates, the dissimilarities caused by stochastic effects remain high even if the template amount and number of PCR cycles was tightly controlled (Figure 7: Jaccard: ca. 10–40%; Bray-Curtis: ca. 5–20%). The variance was not minimized significantly by any of the tested protocols (GLMM: all explanatory variables p-value>0.05, variance explained by maximal model <5%; Information-theoretic approach: only the null model has ΔAICc<2) although the dissimilarities tended to be lower with a smaller number of PCR cycles and higher amount of DNA template.

**Figure 7:**
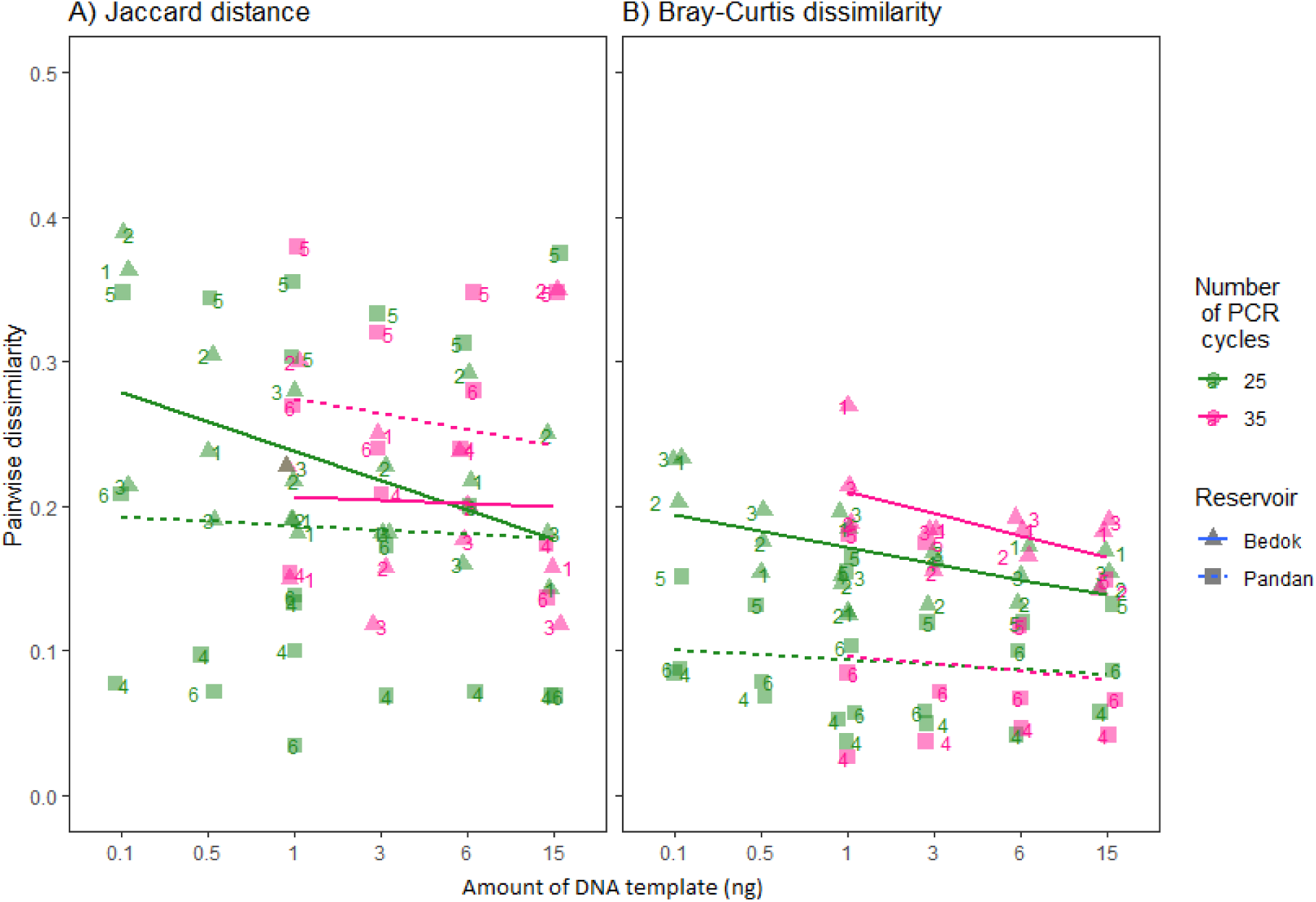
Pairwise dissimilarity (or error) between technical replicates shows no evidence that changes in the number of PCR cycles or in the amount of DNA template resulted in lower dissimilarity (GLM analysis showed that R^2^ of maximal model is low: Jaccard=0.0589; Bray-Curtis=0.0581). Water samples are labeled 1 to 6, each datapoint shows the dissimilarity between the pair of technical replicates subjected to the same PCR protocols. Pairwise dissimilarity was calculated using Jaccard distances (presence/absence data) and Bray Curtis dissimilarity (relative read counts).

As expected, biological replicates within the same reservoir have a wider range of pairwise dissimilarities than technical replicates [biological replicates in Figure 8: Jaccard (0– 50%), Bray-Curtis (ca. 10–40%); technical replicates in Figure 7: Jaccard: (ca. 10–40%); Bray-Curtis: (ca. 5–20%)]. Across technical and biological replicates, dissimilarity measures using relative read counts (Bray-Curtis) yielded slightly more repeatable results than Jaccard. However, the overall reservoir signal remains strong regardless of whether read abundances were used as illustrated by NMDS plots (Figure 6).

**Figure 8:**
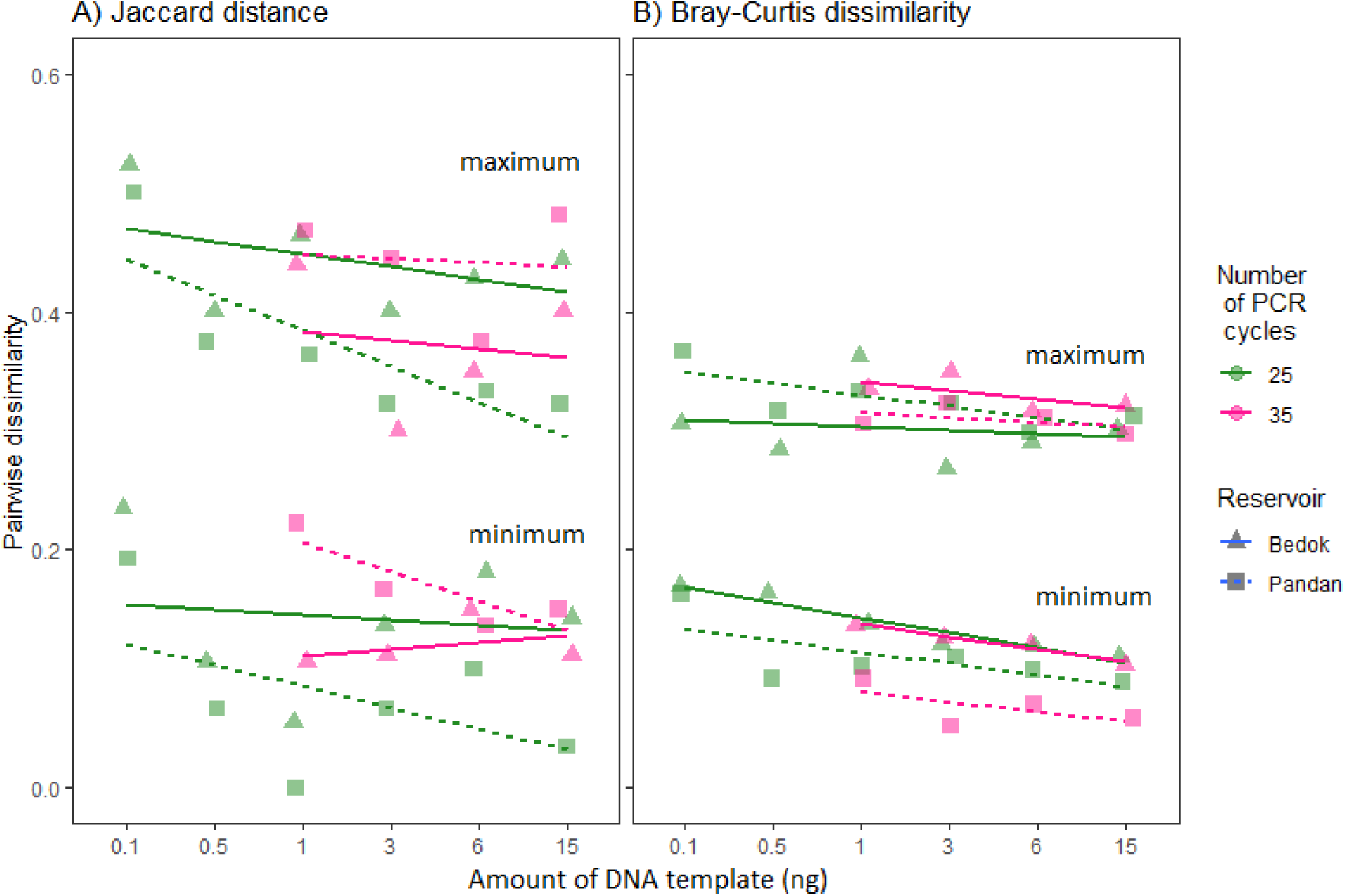
The level of pairwise dissimilarities (or error) between biological replicates spans a wide range (Jaccard: ca. 0–0.5. Bray-Curtis: ca. 0.1–0.4). Only the minimum and maximum pairwise dissimilarity is shown on the graph, although N=6 for each set of molecular conditions and reservoir. Pairwise dissimilarity was calculated using Jaccard distances (presence/absence data) and Bray Curtis dissimilarity (relative read counts).

## 4. Discussion

We here attempted to identify molecular procedures that are suitable for distinguishing two reservoirs based on the eDNA obtained from 15 mL of water. We find that the eDNA signal in the reservoirs is so strong that all laboratory conditions we tested were capable of assigning small samples of water to a particular reservoir. Clews and colleagues (2014), who designed biotic indices for Singapore’s reservoirs, showed that Singapore’s reservoirs were primarily differentiated by trophic status (or productivity), defined in terms of turbidity levels and chlorophyll *a* concentration. Pandan Reservoir and Bedok Reservoir were clearly differentiated by that metric with Pandan Reservoir having higher levels of productivity (see also, Sim et al., 2021), which may explain the higher concentrations of eDNA. Interestingly, Pandan water also had a higher proportion of metazoan reads, despite standardizing molecular conditions and pooling of PCR products in equal volumes (Figure 3). A potential reason is that DNA degradation rates are higher in oligotrophic than eutrophic lakes, as demonstrated in a controlled experiment of lake water (Eichmiller, Best, et al., 2016).

The macroinvertebrate assemblages of the two reservoirs are known to differ with regard to the taxonomic composition (Clews et al., 2014), but we are here able to confirm the differences also for a different set of taxa based on a COI primer designed for Metazoa. Only 28.8% of the 59 mOTUs were shared between reservoirs (Table S2 in Supplementary Information). The mOTUs detected with eDNA were mostly zooplankton and unidentified mOTUs, which, however, are unlikely macroinvertebrates because many of Singaporés macroinvertebrate species have been DNA barcoded (Meier, unpublished). Our findings highlight once more that bioassessment based on eDNA and macroinvertebrate samples are based on different taxa (Gleason et al., 2020; Hajibabaei et al., 2019). We here only amplified metazoan DNA, but the taxonomic signal obtained from the same eDNA samples could surely be multiplied if different primers and genes were used to target different taxa. This implies that 15 mL collected with a remote vehicle has the potential to be more than sufficient for routine bioassessment.

Serial dilution of DNA template amounting to as low as 0.5 ng did not significantly affect the repeatability of results. This implies that PCR template amounts do not have to controlled tightly as long as they remain above this threshold. MOTU compositions were similar for technical replicates across the range of 0.5 ng to 15 ng DNA template amounts (Figures 4, 7). This is congruent with previous studies (Krehenwinkel et al., 2017; Wu et al., 2010) and a convenient finding because much time and cost can be saved that would otherwise be invested in quantifying, diluting and standardizing DNA template amounts. What is arguably less convenient is that the use of equimolar amounts of DNA template and the pooling of PCR products at equal volumes did not prevent higher read coverage for Pandan vs. Bedok Reservoir samples (Figure 3). This problem was further amplified by a larger proportion of metazoan reads in the Pandan data. Overall, we thus conclude that sample adjustments are best made at the read count level, which is increasingly feasible given that sequencing cost continues to decline. With regard to DNA amounts, we only recommend that very low DNA template amounts (0.1 ng) should be avoided because they significantly depress the number of detected mOTUs (Figure 5). There is also some evidence that very low template amounts increases variability (Figure 7; although its effects are not significant). This could generate differences in mOTU profiles (see heatmaps in Figure 4).

If 0.5 ng of DNA were to be used as a benchmark, the minimum volume of water that would need to be sampled and processed can be estimated. In the present study, only 0.681 mL and 0.234 mL of raw water are needed to obtain 1.5 ng of extracted DNA (0.5 ng × 3 PCR replicates) for those sampling sites in Bedok Reservoir and Pandan Reservoir that had the lowest eDNA concentration. Note, however, that this calculation assumes that diluting PCR template amounts is equivalent to reducing the volume of raw water that is DNA extracted. This assumption is known to be violated for soil samples (Ranjard et al., 2003), but we also doubt that anyone would want to sample <1 mL of water; i.e., our main point is that very small volumes of water suffice for bioassessment purposes.

Our present study also suggests that the benefits of using 25 instead of 35 PCR cycles are only slight given that they do not improve repeatability significantly. However, data obtained with different cycling conditions should not be compared. There is some evidence that 35 cycles instead of 25 renders the results obtained from technical replicates less repeatable for Bedok and Pandan samples (Figure 7), but the differences are not statistically significant. On the other hand, comparing mOTU compositions based on data obtained with different cycling conditions may lead to misleading biological conclusions. The main reason is that the use of 35 instead of 25 PCR cycles reduces significantly the number of detected mOTUs (Figure 5). This finding is consistent with the literature [e.g., Kelly et al., (2019)]. In our study, mOTU richness drops because mOTUs that were detected at low read abundance with 25-cycles disappear when the data obtained with 35 PCR cycles were analyzed (Figure 4). This is supported by previous studies suggesting that random subsampling at various steps in molecular protocols affects the eventual read abundance and recovery of rare mOTUs (Kebschull & Zador, 2015; Krehenwinkel et al., 2018; Leray & Knowlton, 2017; Shirazi et al., 2021; Zhou et al., 2011). Random sampling can occur for example, during library preparation (e.g., adaptor ligation) and/or sequencing (e.g., flow-cell ligation) and increasing PCR cycles only increases stochastic effects. In a previous study, Piñol et al. (2015) also demonstrated that increasing PCR cycles reduced the relative read counts of DNA signatures that were amplifying poorly in mock samples. Such signatures are also less likely to survive the bioinformatic filters.

Our study also compared the dissimilarity of mOTUs obtained from technical (=same water sample) and biological replicates (different sites in the same reservoir). Not surprisingly, biological replicates have higher dissimilarities than technical ones (Figures 7, 8), but it was comforting that the site differences were still small compared to the reservoir differences (Figure 6). What was less comforting was the high dissimilarity between technical replicates (Figure 7: Jaccard, presence/absence data: ca. 10–40%; Bray-Curtis, relative read counts: ca. 5–20%). In a way, they measure the inherent error that is produced by a combination of laboratory work and bioinformatic processing. Overall, it would be desirable if error and error variance were lower. For our study, we found that dissimilarity measures based on species and read counts were slightly more repeatable than those based on presence/absence data. Using relative read counts in our comparisons lowered the pairwise dissimilarity between technical replicates. This suggest that using read abundance data is an option for bioassessment.

The conclusions presented here are not without caveats. Firstly, the experiment was conducted on two reservoirs only. It is difficult to tell if the eDNA signals are strong enough to detect subtle differences between other reservoirs, such that the information will be useful for their monitoring and management. These questions are currently being addressed by applying these techniques to a wider range of reservoirs with smaller differences in water quality. Secondly, even fewer of PCR cycles should be tested because they may further reduce stochasticity and lead to more repeatable results.

## Conclusions

Our study suggests that (1) water volumes as low as 1 mL are sufficient to distinguish reservoirs with different water qualities reliably based on Metazoan eDNA signal only. (2) Controlling the amount of PCR template does not improve the repeatability of results as long the template amount exceeds 0.5 µl. (3) Lower PCR cycle numbers (25 vs 35) do not change the repeatability of the results between technical replicates significantly, but higher PCR cycling numbers reduce the number mOTUs that can be used to detect reservoir-signals. Lastly, dissimilarity measures using read abundance data are more reproducible than binary data and may thus be more useful for biotic indices. Our recommendations are also relevant to other applications that require repeatable and reliable community detection.

## Acknowledgements

We thank PUB, Singapore’s National Water Agency, and PUB officers for assisting us in field sampling, especially for giving us access to boats and boat operators. We also thank Jonathan Ho as well as other staff and students of the National University of Singapore (NUS) Evolutionary Biology Lab, and Freshwater and Invasion Biology Lab, for assistance with field sampling. We acknowledge financial support from the Lady Yuen Peng McNeice Graduate Fellowship (awarded to R. K. L.).

## Author Contributions

R. M. and R. K. L. conceived and designed the study. R. K. L. collected the samples and performed the laboratory experiments. R. K. L., R. M. and S. N. K. analyzed the data and interpreted the results. R. K. L., R. M., S. N. K. and D. C. J. Y. wrote the manuscript. All authors have read the approved the final manuscript.

