## Supplementary Materials for "Towards environmental DNA-based bioassessment of freshwater reservoirs with small volumes of water: robust molecular protocols"

### Supplementary Information

**Table S1**: Sampling details, including sampling site, date and exact GPS coordinates.

| **Reservoir** | **Sampling site** | **Sampling date** | **GPS coordinates** |
| --- | --- | --- | --- |
| Bedok | 1 | 26 January 2018 | 01.34066°N, 103.92108°E |
| Bedok | 2 | 26 January 2018 | 01.33888°N, 103.93061°E |
| Bedok | 3 | 26 January 2018 | 01.34497°N, 103.92702°E |
| Pandan | 1 | 24 January 2018 | 01.31203°N, 103.73579°E |
| Pandan | 2 | 24 January 2018 | 01.30378°N, 103.74453°E |
| Pandan | 3 | 24 January 2018 | 01.31813°N, 103.74591°E |

**Table S2**: Few shared metazoan species between reservoirs (29%: 17 out of 59). Most mOTUs with high identities are zooplankton (≥97% similarity) with the majority of mOTUs remaining unidentified.

| **Higher taxonomic identity** | **Family** | **Number of mOTUs** | | | |
| --- | --- | --- | --- | --- | --- |
|  |  | Bedok Reservoir | Pandan Reservoir | Shared | Total |
| Cladocera | Sididae | 1 | 0 | 0 | 1 |
|  | Unidentified | 1 | 0 | 0 | 1 |
| Copepoda | Cyclopidae | 2 | 3 | 2 | 3 |
|  | Unidentified | 1 | 1 | 1 | 1 |
| Oligochaeta | Unidentified | 0 | 1 | 0 | 1 |
| Rotifera | Brachionidae | 1 | 1 | 1 | 1 |
|  | Synchaetidae | 1 | 1 | 0 | 2 |
|  | Trichocercidae | 0 | 3 | 0 | 3 |
|  | Unidentified | 7 | 11 | 4 | 14 |
| Zooplankton | Unidentified | 4 | 2 | 0 | 6 |
| Unidentified | Unidentified | 21 | 14 | 9 | 26 |
|  | **Total** | **39** | **37** | **17** | **59** |

#### 2.6.2 Supplementary Materials and Methods

*DNA extraction: Phenol Chloroform extraction*

Frozen sample mixtures of reservoir water, ethanol and sodium acetate were left to thaw overnight at -30 ⁰C. After thawing, sample mixtures were separated into smaller 50 mL volumes in falcon tubes and spun down (5,500 rcf at 6 ⁰C) for 45 min to pellet eDNA. Supernatant was decanted. From here, DNA pellets were kept separate and processed separately until the end of extraction. DNA pellets were resuspended in 900 mL of cetyltrimethylammonium bromide (CTAB) buffer (0.1 M Tris pH 8; 1.4 M NaCl; 0.02 M EDTA; 20 g/L CTAB) and 20 µL of 1 mg/mL proteinase K buffer (Invitrogen), and transferred into 1.5 mL Eppendorf tubes. Tubes were incubated for 3 hrs at 55 ⁰C.

Following incubation, 600 µL of 25:24:1 phenol:chloroform:isoamyl alcohol (PCI; Biozol) was added to each tube and spun down (17,900 rcf) for 10 min. 600 µL of the aqueous fraction was transferred into a new Eppendorf tube, topped up with 400 µL of fresh PCI and spun down again (17,900 rcf, 10 min). 400 µL of the aqueous fraction was transferred into a new Eppendorf tube and topped up with absolute ethanol 1.5 mL mark. DNA was left to precipitate overnight at -30 ⁰C. Samples were spun down (17,900 rcf at 6 ⁰C) for 30 min. DNA pellets were resuspended in 500 µL of 70% ethanol (with mixing) and spun down again (17,900 rcf) for 15 min. Supernatant was decanted, purified DNA pellets were air-dried, dissolved in molecular-grade water and stored at -30⁰C.

*PCR mixtures and conditions*

PCR mixtures (25 µL total volume) consist of 1X BioReady rTaq buffer (Bioer), 0.2 mM dNTPs (Bioer), 0.2 μg/μL BSA (ACROS Organics), 8 nM of forward and 8 nM of reverse primer (Integrated DNA Technologies), 2U BioReady rTaq DNA Polymerase (Bioer) and 5 μL DNA extract (diluted to desired concentration).

Cycling conditions were as follows: a starting denaturation step of 3 min at 95 ⁰C, followed by 25 or 35 cycles of 30s at 95 ⁰C, 1 min at 45 ⁰C and 30s at 72 ⁰C, and a final extension step of 1 min at 72 ⁰C.

*List of candidate models assessed using the information-theoretic approach (1)*

glm(Dist~cycles*I(DNA^2),data,family=binomial)

glm(Dist~cycles+I(DNA^2),data,family=binomial)

glm(Dist~cycles+DNA+I(DNA^2),data,family=binomial)

glm(Dist~cycles*DNA,data,family=binomial)

glm(Dist~cycles+DNA,data,family=binomial)

glm(Dist~DNA,data,family=binomial)

glm(Dist~cycles,data,family=binomial)

glm(Dist~1,data,family=binomial)

Response variable “Dist” represents either Jaccard distance (binary) or Bray-Curtis dissimilarity (relative read counts) between technical replicates. Both dissimilarity measures were separately assessed. Explanatory variable “cycles” refers the number of PCR cycles, and “DNA” refers to the amount of DNA template used during PCR. The fixed effects include a quadratic term for DNA in some models (I(DNA^2)). Data were fitted with binomial GLM, using the glm function as implemented in the R package *lme4* v1.1-27.

*List of candidate models assessed using the information-theoretic approach (2)*

glmer(mOTU~cycles*I(DNA^2)+(1|Res),data,nAGQ=1,family=poisson)

glmer(mOTU~cycles+I(DNA^2)+(1|Res),data,nAGQ=1,family=poisson)

glmer(mOTU~cycles+DNA+I(DNA^2)+(1|Res),data,nAGQ=1,family=poisson)

glmer(mOTU~cycles*DNA+(1|Res),data,nAGQ=1,family=poisson)

glmer(mOTU~cycles+DNA+(1|Res),data,nAGQ=1,family=poisson)

glmer(mOTU~DNA+(1|Res),data,nAGQ=1,family=poisson)

glmer(mOTU~cycles+(1|Res),data,nAGQ=1,family=poisson)

glmer(mOTU~1+(1|Res),data,nAGQ=1,family=poisson)

Response variable “mOTU” represents the number of mOTUs detected, or mOTU richness. Explanatory variable “cycles” refers the number of PCR cycles, and “DNA” refers to the amount of DNA template used during PCR. The fixed effects includes a quadratic term for DNA in some models (I(DNA^2)). The random effect “(1|Res)” controls for the differences between reservoir sampled. Data were fitted with poisson GLMMs and Laplace approximation, using the glmer function as implemented in the R package *lme4* v1.1-27.
